# Circadian time series proteomics reveals daily dynamics in cartilage physiology

**DOI:** 10.1101/654855

**Authors:** Michal Dudek, Constanza Angelucci, Jayalath P.D. Ruckshanthi, Ping Wang, Venkatesh Mallikarjun, Craig Lawless, Joe Swift, Karl E. Kadler, Judith A. Hoyland, Shireen R. Lamande, John F. Bateman, Qing-Jun Meng

## Abstract

**Objectives:** Articular cartilage undergoes cyclical heavy loading and low load recovery during the 24-hour day/night cycle. We investigated the daily changes of protein abundance in mouse femoral head articular cartilage by performing 24-hour time-series proteomics study.

**Methods:** Tandem mass spectrometry analysis was used to quantify proteins extracted from mouse cartilage. Bioinformatics analysis was performed to quantify rhythmic changes in protein abundance. Primary chondrocytes were isolated and cultured for independent validation of selected rhythmic proteins.

**Results:** 145 rhythmic proteins were detected. Among these were key cartilage molecules including CCN2, MATN1, PAI-1 and PLOD1 & 2. Pathway analysis revealed that proteins related to protein synthesis, cytoskeleton and glucose metabolism exhibited time-of-day dependent peaks in their abundance. Meta-analysis of published proteomics datasets from articular cartilage revealed that numerous rhythmic proteins were dysregulated in osteoarthritis and/or ageing.

**Conclusions:** Our circadian proteomics study revealed that articular cartilage is a much more dynamic tissue than previously thought. Chondrocytes exhibit circadian rhythms not only in gene expression but also in protein abundance. Our results clearly call for the consideration of circadian timing in understanding cartilage biology, osteoarthritis pathogenesis, treatment strategies and biomarker detection.

## 1. Introduction

Articular cartilage (AC) is a thin connective tissue that covers the ends of long bones and is critical for the smooth, frictionless movement of joints. The importance of AC in the joint is exemplified in osteo- and rheumatoid arthritis in which loss of cartilage causes joint space narrowing and pain, leading to immobilisation and long-term disability. Time-of-day variation in symptoms – including pain, swelling and joint stiffness – has been reported in the clinical literature and diurnal changes in serum biomarkers (e.g. COMP and IL-6) for arthritic patients have been reported [1]. However, a molecular understanding of the time-of-day variation in AC physiology has been lacking. Recent studies in mice have shown that AC contains intrinsic circadian clocks driving 24-hour rhythmic expression of hundreds of genes. How rhythmic gene expression leads to diurnal changes in cartilage physiology warrants further investigation.

Cycles of day and night govern the circadian (‘about a day’) rhythms of rest and activity, physiology and metabolism of most animals on the surfaces of our planet. In mammals, a central circadian clock in the anterior hypothalamus of the brain co-ordinates the rhythmic behaviour (such as the sleep/wake cycle), and synchronizes various local oscillators that are found in most peripheral organs. At the molecular level, the circadian oscillator is based on a transcriptional/translational feedback loop composed of interlocked transcriptional activators (BMAL1 and Clock) and repressors (CRYs and PERs). In addition to driving the expression of core clock genes, these clock factors control rhythmic expression of genes in a tissue specific manner [2]. Recently, work from our group and others have demonstrated autonomous circadian clocks in chondrocytes within the articular cartilage. The cartilage circadian rhythm becomes disrupted in mouse OA models and human OA, linking circadian timing mechanism to the maintenance of homeostasis and pathogenesis of cartilage [3 4]. Crucially, with the refinement of mass spectrometry technology, there are increasing numbers of studies comparing protein abundance in healthy and arthritic cartilage, as well as studies attempting osteoarthritis biomarker discovery. However, none of these studies have taken into account the time-of-day when the cartilage was harvested and processed.

In this study, we performed a circadian time-series proteomics analysis to reveal the extent of rhythmic proteins in femoral head articular cartilage from mice kept under a 12 hour/12 hour light/dark cycle. Our results revealed daily 24-hour dynamics of 145 proteins in cartilage, including extracellular matrix (ECM) and ECM-related proteins. These results reinforce circadian timing as a new dimension in understanding cartilage function and disease, and call for the consideration of time-of-day as a critical factor in experimental design, data interpretation, standardisation of biomarker detection and timing of therapeutic interventions in joint diseases.

## 2. Methods

### 2.1 Preparation of cartilage samples

All animal studies were performed in accordance with the 1986 UK Home Office Animal Procedures Act. Approval was provided by the Animal Welfare Ethical Review Board (AWERB) of the University of Manchester (approval no. 50/2506). Mice were maintained at 20-22°C, with standard rodent chow available ad libitum and under 12:12 hr light dark schedule (light on at 6 am; light off at 6 pm). 72 male BALB\c mice were sacrificed at 2 months of age. Samples were taken from 6 mice per time-point every 4 hour for 48 hours. Articular cartilage from the head of the proximal femur was separated from the subchondral bone as described previously [3]. Cartilages from two hips of one mouse were pooled together and washed 3 times in PBS with protease (Roche 11836170001) and phosphatase inhibitors (Sigma P0044 and P5726) and subsequently snap frozen in liquid nitrogen.

### 2.2 Protein extraction

Cartilage tissues were pulverized using a liquid-nitrogen-cooled tissue grinder and proteins extracted as previously described [5]. Briefly, cartilage samples were reconstituted in 100 μL of 100 mM Tris acetate buffer pH 8.0 containing 10 mM EDTA and protease/phosphatase inhibitors and deglycosylated by treatment with 0.1 units of chondroitinase ABC for 6 h at 37 °C. Proteins were sequentially extracted in a chaotropic buffer containing guanidine hydrochloride (4M GuHCl, 65 mM DTT, 10 mM EDTA in 50 mM sodium acetate, pH 5.8). Protein samples were precipitated with nine volumes of ethanol, washed once in 70% ethanol, then resuspended in 120 μL of solubilisation buffer (7 M urea, 2 M thiourea, and 30 mM Tris, pH 8.0) and the volume was adjusted to achieve a concentration of ∼1 mg/mL, as estimated using the EZQ protein quantitation protocol (Thermo Fisher). Samples were then stored at −80 °C until required. Protein samples were analysed by SDS-PAGE and detected by silver staining as previously described.

### 2.3 Peptide sample preparation and analysis by Nanoliquid Chromatography and LTQ-Orbitrap Tandem Mass Spectrometry

Protein samples for LC-MS/MS analysis were sequentially reduced and alkylated under nitrogen by incubation in 10 mM dithiothreitol (overnight at 4 °C) then 50 mM iodoacetamide (2 h at 25 °C in the dark). Proteins were co-precipitated with 1 µg trypsin (Promega) overnight at −20 °C in 1 ml methanol. The trypsin-protein precipitates were washed once with chilled methanol, dried and reconstituted in 100 mM ammonium bicarbonate, followed by trypsinization at 37 °C for 5 h, with addition of 1 µg trypsin after 2 h. Digests were terminated by freezing on dry ice. Samples were dissolved in of 0.1% formic acid, 3% acetonitrile and applied to 30K cutoff spin filter column (Millipore Ultracel YM-30). Mass spectrometry was performed by the Mass Spectrometry and Proteomics Facility (Bio21 Molecular Science and Biotechnology Institute, University of Melbourne). LC-MSMS was carried out on a LTQ Orbitrap Elite (Thermo Scientific) with a nanoelectrospray interface coupled to an Ultimate 3000 RSLC nanosystem (Dionex). The nanoLC system was equipped with an Acclaim Pepmap nano-trap column (Dionex – C18, 100 Å, 75 µm × 2 cm) and an Acclaim Pepmap analytical column (Dionex C18, 2µm, 100 Å, 75 µm × 15 cm). 2 µl of the peptide mix was loaded onto the trap column at an isocratic flow of 5 µl/min of 3% CH3CN containing 0.1% formic acid for 5 min before the enrichment column is switched in-line with the analytical column. The eluents used for the liquid chromatography were 0.1% (v/v) formic acid (solvent A) and 100% CH3CN/0.1% formic acid (v/v) (solvent B). The flow following gradient was used : 6% to 10% B for 12 min, 10% to 30% B in 20 min, 30% to 45% B in 2 min, 45% to 80% in 2 min and maintained at 80% B for 3 min followed by equilibration at 3% B for 7min before the next sample injection. The LTQ Orbitrap Elite mass spectrometer was operated in the data dependent mode with nano ESI spray voltage of +2.0 kv, capillary temperature of 250°C and S-lens RF value of 60%. A data dependent mode whereby spectra were acquired first in positive mode with full scan scanning from m/z 300-1650 in the FT mode at 240,000 resolution followed by Collision induced dissociation (CID) in the linear ion trap with ten most intense peptide ions with charge states ≥2 isolated and fragmented using normalized collision energy of 35 and activation Q of 0.25.

### 2.4 Bioinformatic analysis

Maxquant version 1.5.8.3 [6] was used to analyse the raw files from the LTQ Orbitrap Elite (Thermo) with ultimate 3000 nanoLC. Spectra were searched against a Fasta file of the complete *Mus musculus* proteome downloaded from Uniprot, using Maxquant internal search engine Andromeda. Settings in Maxquant for Label free quantification were left as default except that ‘match between runs’ was selected with a match time window of 2mins. Specific enzyme was Trypsin/P with max missed cleavages set to 2. Peptide and protein false discovery rates (FDR) were set to 0.01 with maximal posterior error probability (PEP) set to 0.01. The minimal peptide length was set to 7 and minimum peptide and ‘razor+ unique peptides’ was set to 1. Unique and razor peptides were used for quantification as recommended by Cox *et al*. [7] with a minimum ratio count of 2. Normalised intensity values (LFQ intensity) was used for quantification and the protein groups results file was analyse using Perseus (1.5.3.1). Reverse and ‘only identified by site’ hits were removed. LFQ intensity data was log transformed. Intensity data in protein groups were further required to be detected in half of the total number of samples. We then use meta2d, a function of the R package Metacycle [8], to evaluate periodicity in the proteomic data. Rhythmic genes were then clustered with cluster and Rtsne of the R package. Rose plots were generated from the proteome and using metad2_p-value 0.05 threshold. Principal component analysis (PCA) was performed by singular value decomposition on rhythmic proteins (metad2_p-value<0.05) after z-score normalisation. Protein interaction network was generated in CytoScape [9] using the STRING plugin [10]. Only experimentally determined interactions from curated databases were used. Nodes without interaction partners were removed from the graph. Extracellular Matrix proteins were determined by comparison with the matrisome database (matrisome.org) [11].

### 2.5 Mouse primary chondrocyte culture

Primary chondrocytes from 5 day old mice were isolated according to published protocol [12]. Briefly, 5 day old C57bl/6 PER2::Luc mice were sacrificed by decapitation. Knee, hip and shoulder joints were dissected, and any soft tissue removed. Joint cartilage subjected to pre-digestion with collagenase D 3mg/mL in DMEM two times for 30 min at 37°C with intermittent vortex to remove soft tissue leftovers. Subsequently, the cartilage was diced using a scalpel and digested overnight at 37°C. Cells were dispersed by pipetting and passed through 70 µM cell strainer. Cell suspension was then centrifuged and the pellet was re-suspended in DMEM/F12 with 10% FBS and plated in T75 flasks. Cells were passaged only once before performing experiments.

### 2.6 Western Blotting

Western blotting was performed according to standard procedures. Primary antibodies for CTGF (Abcam ab6992), PAI-1 (Abcam ab66705), MATN1 (Abcam ab106384) and alpha tubulin (Sigma T9026) were used in 1:2000 dilution. Secondary antibodies (LI-COR IRDye 800CW and 680RD) were used in 1:20000 dilution. WB was quantified using the LI-COR Odyssey Imaging System.

## 3. Results

To identify rhythmic changes in the proteome, we performed a 48-hour time series proteomic analysis of femoral head articular cartilage using young mice (8 weeks of age) collected at 4 hour intervals across two 12-hour day/12-hour night cycles. We were able to identify 1177 proteins by a minimum of two unique peptides (Supplementary Table 1). Of these, 743 overlapped with proteins in datasets from two single time point studies using similar methods [5, 13]. Importantly, the time-series samples allowed us to identify 425 proteins not detected in previous studies, indicating the existence of time-dependently expressed molecules (Supplementary Figure 1A).

**Table 1.**
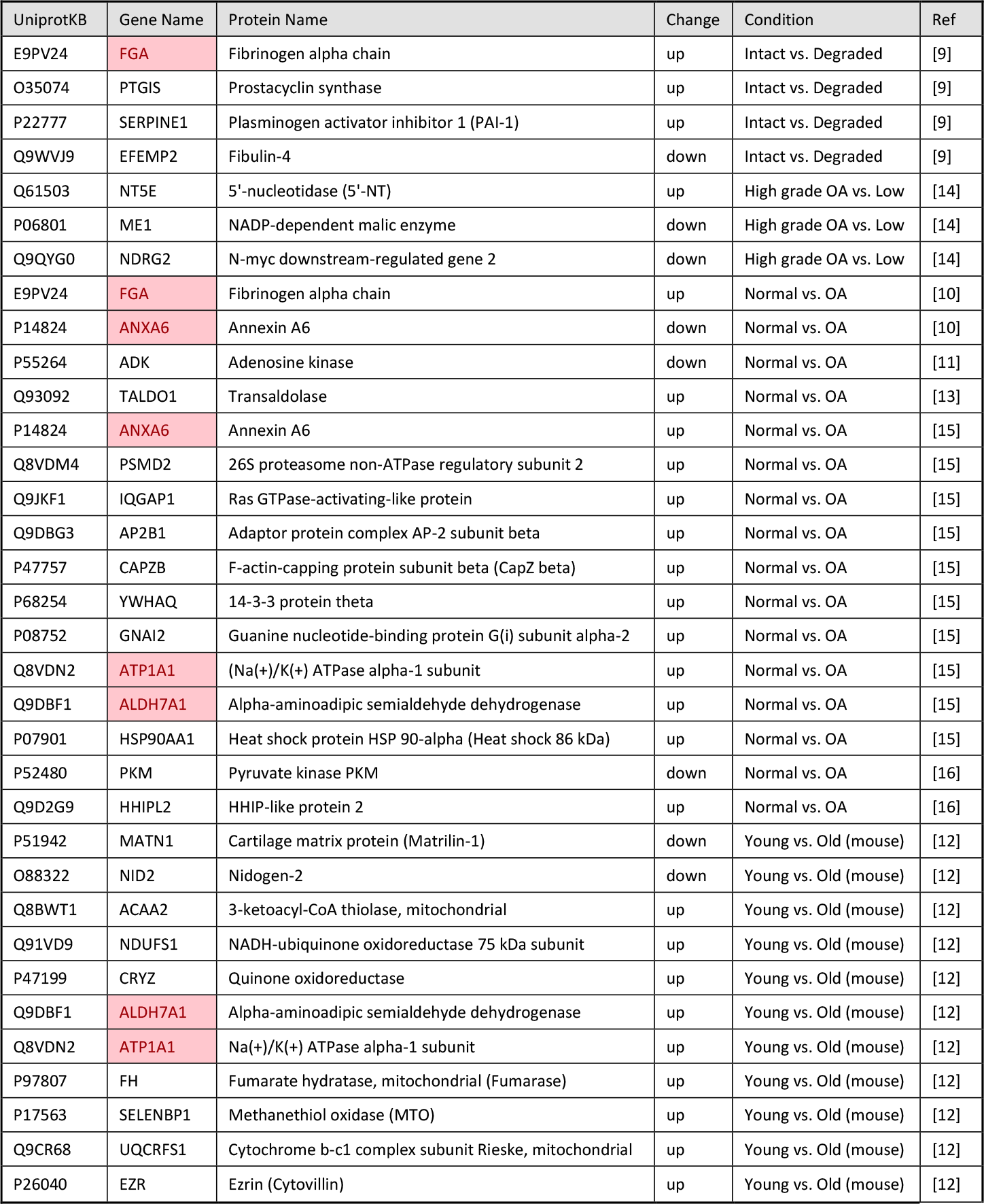
Meta-analysis of published proteomic datasets comparing normal vs. osteoarthritic cartilage and young vs. aged cartilage. Rhythmic proteins dysregulated in more than one study are highlighted in red.

Statistical analysis of the time-series using MetaCycle revealed 145 rhythmic proteins (12.3% of total identified) with integrated p value <0.05 (Figure 1A). Principal component analysis of the samples revealed that the major two components separating the data organised the samples into day and night clusters (Supplementary Figure 1B). Most of the rhythmic proteins peaked during late night at 2-4 am (active phase in mice).

**Figure 1.**
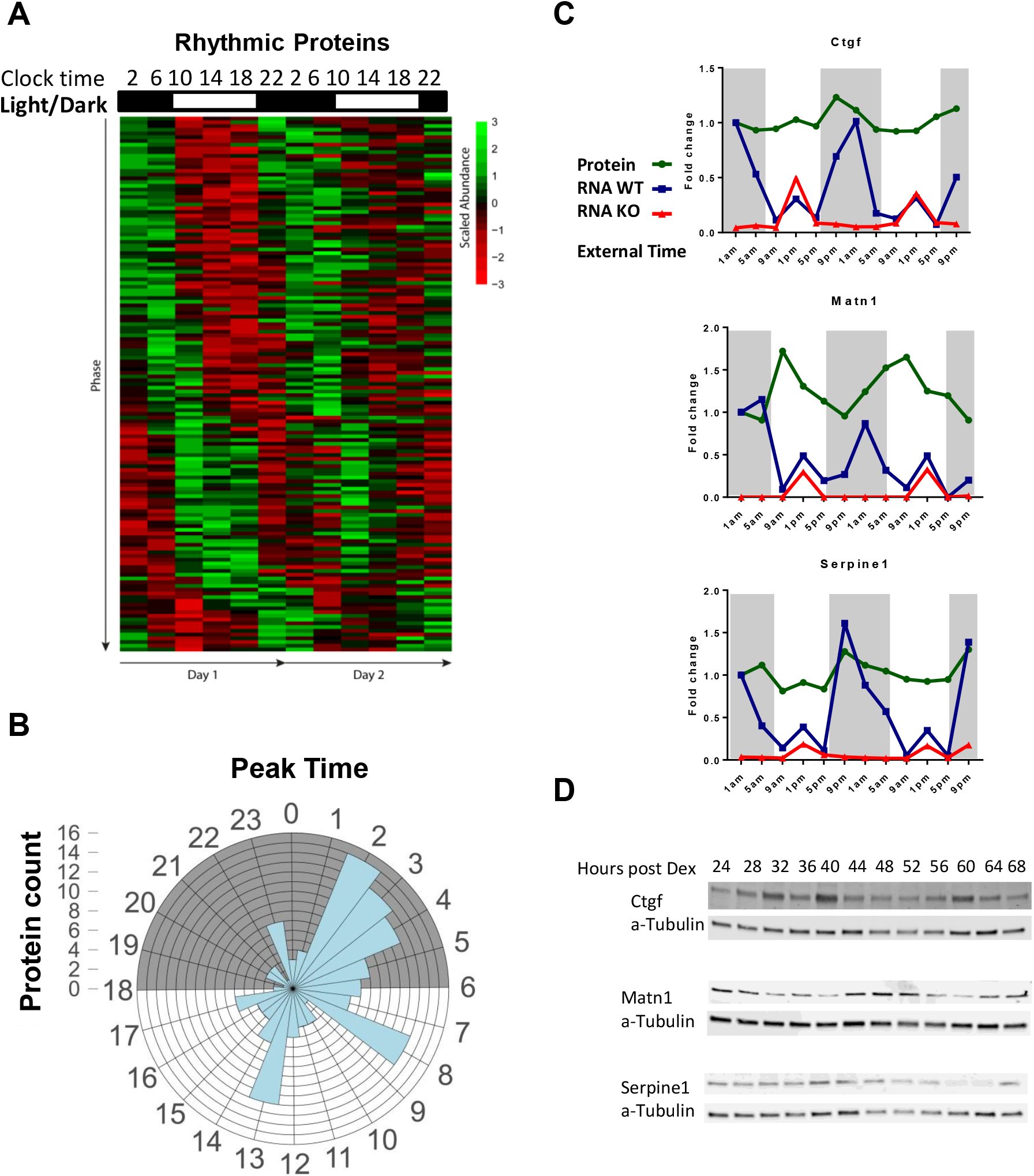
Articular cartilage exhibits circadian rhythm in protein abundance. **A.** Heat map of the 48-hour time-series experiment showing 145 rhythmic proteins identified by mass spectrometry in mouse hip articular cartilage (MetaCycle integrated p value <0.05). **B.** Rose plot showing distribution of peak abundance of rhythmic proteins within the 24-hour circadian cycle. Shaded area indicates the dark phase. **C.** Fold changes of protein abundance (by Mass Spec) and gene expression (by RNAseq) of selected molecules in mouse hip articular cartilage. Expression of these genes is affected by disruption of the circadian clock (Col2a1-Cre/*Bmal1* KO). **D.** Representative results confirming changes in protein abundance over a period of 48 hours by western blotting in primary mouse chondrocyte culture synchronised by dexamethasone.

A number of rhythmic proteins also showed rhythmicity at mRNA levels in our mouse cartilage RNAseq time-series dataset ([3] and unpublished), such as CTGF (CCN2), matrilin 1 and serpine 1(PAI-1). The mRNA expression levels and rhythmic patterns for all three proteins were greatly reduced in the cartilage specific *Bmal1* KO mouse, indicating functioning circadian clock as an essential regulator for their expression (Figure 1D). Daily variations in protein abundance were independently validated in clock-synchronized mouse primary chondrocytes by western blotting (Figure 1E).

STRING interaction analysis of the rhythmic proteins indicate that many proteins that physically interact with each other and take part in the same biological processes largely peak together during the same part of the day (Figure 2A). Proteins responsible for mRNA processing and ribosomal proteins peak in the afternoon, likely to be in preparation for the active phase. Proteins related to ATP synthesis peak in the evening (early active phase) and glucose metabolism peaks late at night (late active phase). Proteins associated with the cytoskeleton peak in the morning (early resting phase). These data indicate temporal segregation of different cellular events in articular cartilage within the 24-hour day.

**Figure 2.**
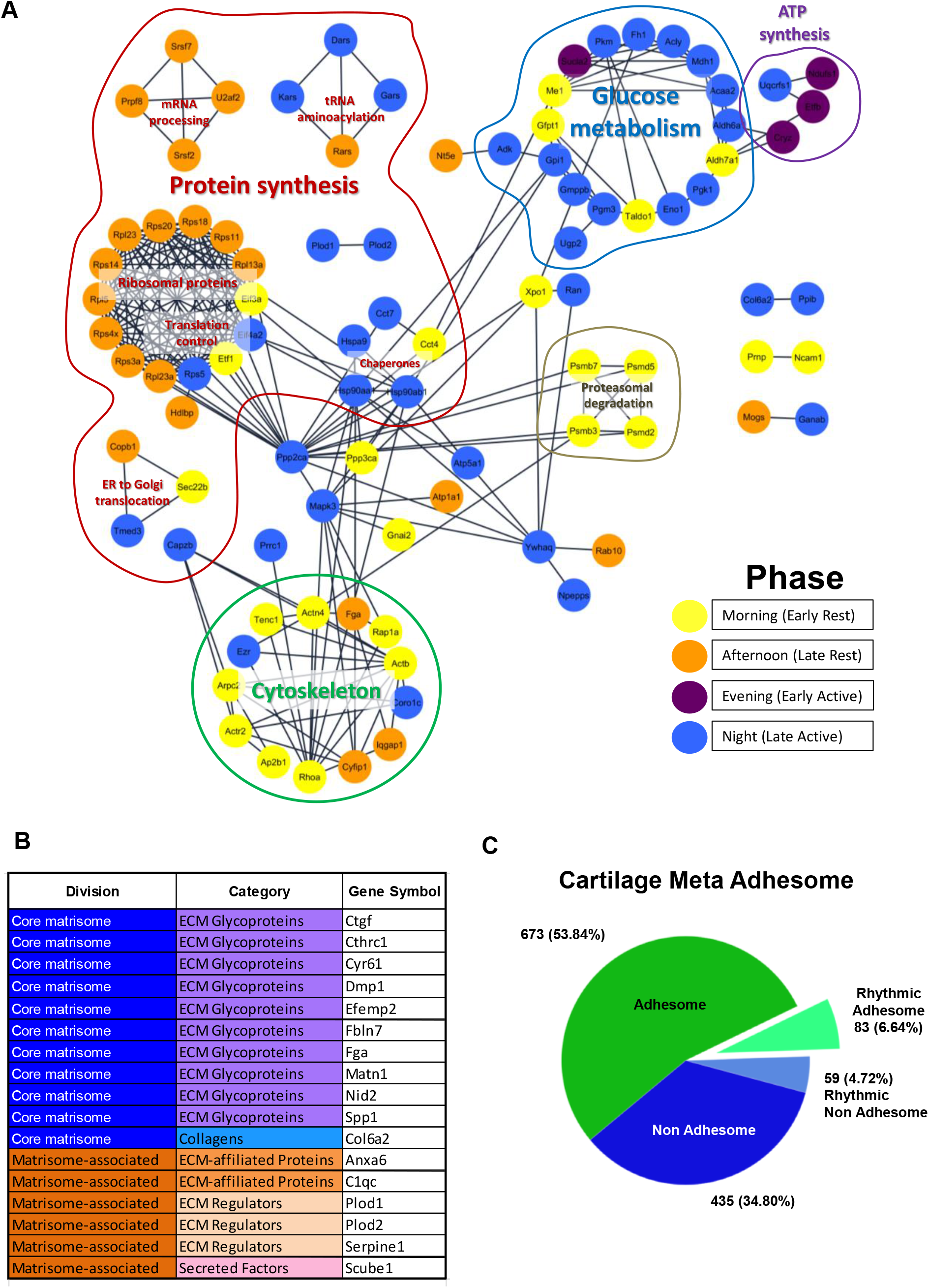
Protein interaction network of rhythmic proteins shows time of day partitioning of cellular processes. **A.** Protein interaction network generated using the STRING plugin for CytoScape utilising interactions from experimental data and curated databases. Nodes are colour coded by peak time of protein abundance. **B.** List of rhythmic extracellular matrix proteins identified as core matrisome or matrisome-associated components [11]. **C.** Pie chart showing proportion of rhythmic adhesion associated proteins as compared with the meta-adhesome.

Although cartilage ECM proteins are thought to berelatively stable, when the rhythmic proteomics dataset wascompared with the matrisome database (http://matrisome.org/) [11], we identified 17 rhythmic ECM proteins including CTGF (CCN2), CYR61 (CCN1), DMP1, Osteopontin, Plod1 and Plod2 which play key regulatory functions in cartilage (Figure 2B). For instance, Plod1 and 2 are responsible for hydroxylation of lysine during collagen synthesis, supporting a role of circadian rhythm in regulating the post-translational modification of ECM components, including collagens. Our data also revealed 5 rhythmic core adhesome proteins (Actn4, Hp1bp3, Iqgap1, Ppib and Rpl23a) and 83 overlapping with the extended meta-adhesome list [14] (Figure 2C and Supplementary Table 2).

Further comparison with published ageing or osteoarthritis-regulated cartilage proteomic studies [15-22] revealed a list of 34 rhythmic proteins being dysregulated in disease condition and/or in aging (Table 1).

## 4. Discussion

Circadian proteomic studies have been performed in other tissues (e.g. liver, SCN, plasma) [23-25] and revealed profound dynamics in protein abundance and phosphorylation. To the best of our knowledge this is the first study aiming to characterise circadian proteome of the articular cartilage.

The discovery that 12% of all extractable cartilage proteins show a daily 24-hour rhythm indicate that the articular cartilage is a much more dynamic tissue than previously thought, with many proteins being synthesized and/or turned over on a daily basis. Most of the rhythmic proteins peak during late active phase. This indicates possible adaption of chondrocyte function to the 24-hour day, in order to anticipate and respond to bouts of activity and related demand for protein synthesis and tissue repair thereafter. Many of the rhythmic proteins also show rhythmicity at mRNA levels in our RNAseq data. Dysregulation of these genes in *Bmal1* KO cartilage suggests endogenous clock control in contrast to merely responding to daily mechanical loading. This is in line with recent study by Chang *et al*. [26] showing that collagen type I in mouse tail tendon is rhythmically secreted and assembled and that ∼10% of tendon proteome is rhythmic. Consistent with these findings, a large portion of the machinery related to protein synthesis exhibited a circadian pattern in abundance in cartilage. Rhythmic changes in the abundance of adhesion molecules suggest that chondrocytes may change the way they interact with ECM during a 24-hour cycle.

The emerging importance of the circadian rhythm in physiology adds a new dimension to the complexity of biological systems and poses new challenges to understanding disease processes and biomarker discovery. Our meta-analysis of published proteomic datasets revealed over 20% of rhythmic proteins to be dysregulated in osteoarthritis or with aging. Changes in the abundance of these proteins in ageing or osteoarthritic cartilage could be due to loss of clock function or change of circadian phase. Indeed, we have previously shown that IL-1 dampens the clock rhythm in cartilage and that expression of clock components is dysregulated in osteoarthritis [3, 4]. Therefore, caution should be exercised when measuring osteoarthritis biomarkers since some of them may be rhythmic and time of sampling could have a profound effect on results. Finally, understanding of the rhythmic processes occurring in cartilage may allow optimization of existing osteoarthritis therapies, as well as the identification of new approaches for restoring tissue homeostasis, e.g. through resetting of the circadian clock.

## Contributors

Q-JM and MD designed the study and wrote the manuscript. MD, CA, JPDR performed the experiments. MD, CA, PW, VM and CL performed the bioinformatics analysis. MD, CA, JPDR, PW, VM, CL, JS, KEK, JAH, SRL, JFB and Q-JM analysed the data.

## Patient consent for publication

Not required.

## Data sharing statement

Upon acceptance of the manuscript, all annotated data supporting this study will be deposited in public repositories.

---

**Supplementary Figure 1.**
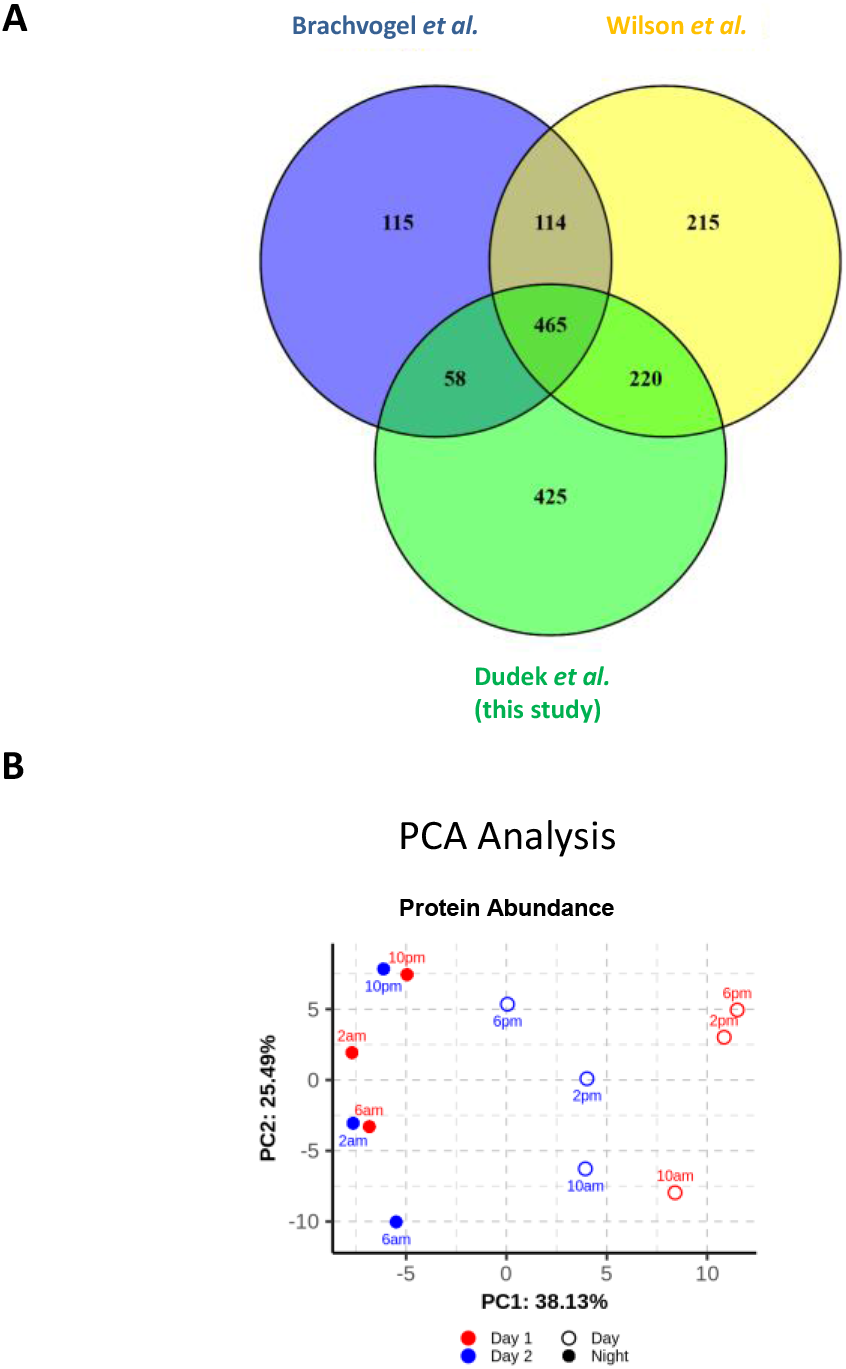
**A.** Venn diagram showing overlap between cartilage proteins identified in the 48-hour time-series experiment and previously published mouse cartilage datasets using similar protein extraction methods [6, 7]. **B.** PCA analysis of proteomic dataset showing temporal separation of the samples depending on the time of day.

